# The biaxial mechanics of thermally denaturing skin - Part I: Experiments

**DOI:** 10.1101/2021.06.04.447116

**Authors:** William D. Meador, Gabriella P. Sugerman, Adrian Buganza Tepole, Manuel K. Rausch

## Abstract

The mechanics of collageneous soft tissues, such as skin, are sensitive to heat. Thus, quantifying and modeling thermo-mechanical coupling of skin is critical to our understanding of skin’s physiology, pathophysiology, as well as its treatment. However, key gaps persist in our knowledge about skin’s coupled thermo-mechanics. Among them, we haven’t quantified the role of skin’s microstructural organization in its response to superphysiological loading. To fill this gap, we conducted a comprehensive set of experiments in which we combined biaxial mechanical testing with histology and two-photon imaging under liquid heat treatment. Among other observations, we found that unconstrained skin, when exposed to high temperatures, shrinks anisotropically with the principle direction of shrinkage being aligned with collagen’s principle orientation. Additionally, we found that when skin is isometrically constrained, it produces significant forces during denaturing that are also anisotropic. Finally, we found that denaturation significantly alters the mechanical behavior of skin. For short exposure times, this alteration is reflected in a reduction of stiffness at high strains. At long exposure times, the tissue softened to a point where it became untestable. We supplemented our findings with confirmation of collagen denaturation in skin via loss of birefringence and second harmonic generation. Finally, we captured all time-, temperature-, and direction-dependent experimental findings in a hypothetical model. Thus, this work fills a fundamental gap in our current understanding of skin thermo-mechanics and will support future developments in thermal injury prevention, thermal injury management, and thermal therapeutics of skin.

## Introduction

All biological tissues, including collageneous soft tissues, are sensitive to heat. In fact, collagen, the predominant structural protein in collageneous soft tissues, denatures at temperatures significantly exceeding body temperature [1]. That is, at temperatures well above 37°C collagen’s triple helix structure collapses as inter- and intra-molecular cross-links are broken, and the resulting unstructured collagen molecules dissociate [2]. Eventual loss of functional collagen transforms these tissues into gel-like, gelatinous materials. This transformation comes with a loss of load-bearing ability, i.e., stiffness and toughness [3]. Additionally, soft tissues shrink significantly during denaturation [4]. These processes are not instantaneous but follow exponential functions that demonstrate Arrhenius-type temperature-dependence [5]. Previously, Chen et al. demonstrated this behavior for chordae tendineae, a highly collageneous cardiovascular soft tissue, and described time-temperature equivalence for this tissue [6]. In a later study, Chen et al. further demonstrated time-temperature-stress equivalence in similar collageneous tissues highlighting the multibiophysical character of collageneous soft tissue denaturation [7]. One-dimensional, or uniaxial tests, where later extended by the same investigators to two-dimensional, or biaxial, studies of membranous materials. Those studies showed that time-, temperature-, and load-dependence may be similarly observed as in above studies but also revealed that uniaxial data cannot be simply extrapolated to multi-axial data [8, 9].

Understanding thermally-induced denaturation of collageneous soft tissues is clearly important from a basic pathophysiological perspective. Additionally, understanding this biophysical process is critical from a therapeutic perspective. For example, Boronyak et al. and Price et al. proposed using thermal denaturation-induced shrinkage to treat myxomatous mitral valve regurgitation [10, 11, 12]. Similarly, thermally-induced tissue shrinkage has been used to treat shoulder instability [13, 14] and to close femoral access wounds [15]. These are just a few of many applications where thermal denaturation of collageneous tissues was applied therapeutically.

For skin specifically, studies of thermal denaturation are critical both from a pathophysiological perspective as well as from a therapeutic perspective. As skin’s function is to protect us from environmental threats, such as heat, understanding and quantifying its response to superphysiological loading, i.e., thermally-induced denaturation, is vital [16, 17]. Similarly, there are skin-specific medical therapies that use thermal loading whose multibiophysical effects and limits should be known for safe use, such as laser-ablation [18]. Finally, aesthetic trends such as branding make direct use of thermal denaturation, again, requiring knowledge of the underlying processes for safe application [19].

Because of skin’s critical role in thermoregulation, there have been numerous studies investigating skin under physiological conditions with and without including effects on or due to mechanics [20, 21]. On the other hand, studies of skin mechanics under superphysiological conditions are sparse. Notable exceptions include those by Zhou et al and Zu et al. for example [22, 23]. They demonstrated that thermally denaturing skin exhibits constitutive behavior that is temperature and load-rate dependent. However, there have been no reports of skin biaxial mechanics of thermally denaturing skin under both isotonic and isometric conditions that considered the tissue’s anisotropy. The objective of our current work is to fill this gap. To this end, we conduct three distinct experiments: i) isotonic, denaturation-induced shrinkage measurements at superphysiological temperatures, ii) isometric, denaturation-induced tension measurements at the same superphysiological temperatures, iii) and finally, biaxial test of skin’s anistropic mechanical properties before and after heat treatment. We supplement and interpret these data by means of classic histology and two-photon microscopy-based structural investigations.

## Materials and Methods

### Skin samples

All animal experiments were conducted in accordance with NIH’s Guide for Care and Use of Laboratory Animals and after approval from our local institutional animal care and use committee. All mice were humanely sacrificed via CO_2_ inhalation before we removed the hair from the dorsal skin regions with clippers and a chemical depilatory agent (Nair, Churd & Dwight, Inc., Ewing, NJ, United States). For the isometric biaxial test samples, we applied an ink stamp with known dimensions (6mm x 6mm) to four dorsal skin regions before excising those regions as 10mm x 10mm square samples. For the isotonic shrinkage and two-photon studies, we used biopsy punches to remove up to eight 6mm dorsal skin samples.

### Isotonic shrinkage experiments

First, we placed discoid skin samples of 6mm diameter on a 1xPBS-wetted, calibrated grid and took photographs of the samples. Next, we submerged those samples in 37°C, 55°C, 75°C, or 95°C 1xPBS for varying durations. After removing the samples from the bath we immediately submerged the tissue in a 37°C 1xPBS bath, i.e., “quenched” the tissue. We then took photographs of those samples on a 1xPBS-wetted calibrated grid again. See Figure 1A for a depiction of the protocol.

**Figure 1:**
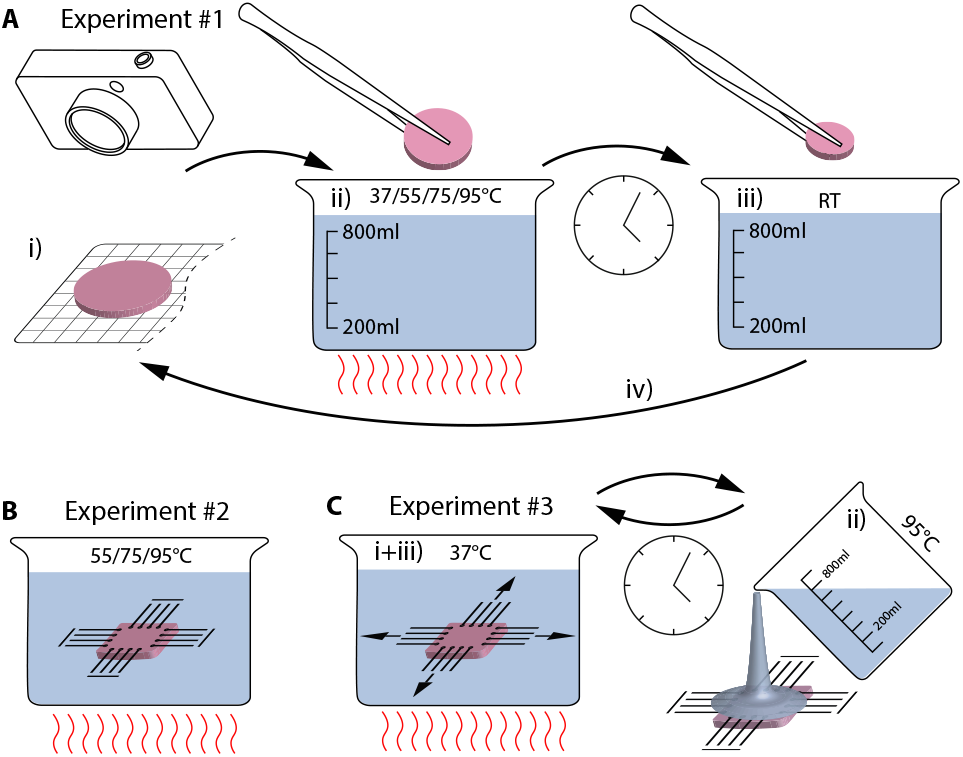
Experimental design of our study. A) Experiment #1 describes our biaxial shrink-age study under traction-free or isotonic boundary conditions in which we i) took images of excised skin samples on a calibrated grid, before ii) submerging the samples in a 1xPBS bath heated to 37°C, 55°C, 75°C, or 95°C for various exposure times. We halted the temperature treatment by iii) submerging samples in 37°C water before, iv) yet, again, taking images of the samples on a calibrated grid. B) Experiment #2 describes our isometric biaxial experiment, in which we i) quantified the biaxial constitutive behavior of excised skin samples in a 37°C water bath, before ii) exchanging the bath with 55°C, 75°C, or 95°C water for 5 minutes during which we measured the resulting forces while fixing the boundaries of the sample in its previously stress-free reference configuration. Subsequently, we iii) retested the biaxial constitutive behavior of the samples. C) Experiment #3 describes our second biaxial experiment, in which we i) quantified the biaxial constitutive behavior of excised skin samples in a 37°C water bath, before ii) removing the water bath and treating the samples with a 90°C water shower for varying exposure times during which we measured the resulting forces while fixing the boundaries of the sample in its previously stress-free reference configuration. Subsequently, we iii) retested the biaxial constitutive behavior of the samples

### Isometric biaxial tension experiments

We mounted the 10mm x 10mm square samples onto our customized biaxial tensile tester (Cell Scale, Biotester, Waterloo, Canada) within 4 hours of excision. Once mounted, we established an approximately stress-free reference configuration by loading the tissue to an equibiaxial preload of 20mN. Next, we conducted our standard biaxial protocol to equibiaxial force of 1000mN while the samples were submerged in 37°C 1xPBS. We repeated this loading cycle 20 times until the material response stabilized. After recording the tissues baseline constitutive response, we conducted one of two protocols. During the first set of experiments, we let the tissue recover from initial biaxial testing for 5 minutes in 37°C 1xPBS. After this recovery period, we replaced the fluid bath with either 55°C, 75°C, or 95°C 1xPBS and recorded the developing force under isometric tension, i.e., with fixed boundary conditions. We recorded force and temperature of all samples for the ensuing 5 minutes until the material response equilibriated. During the second, separate set of experiments, we also let the tissue recover for 5 minutes following the initial biaxial testing. After this period, we lowered the tissue bath so that the samples were suspended on our rake system and then gently poured 90°C 1xPBS for between 2-30 seconds onto the samples before re-submerging the samples in 37°C 1xPBS to abruptly stop the heat treatment. We continuously recorded skin temperature with a thermal imaging camera (FLIR, A35/65, Wilsonville, OR, United States) and force while we poured 1xPBS onto our samples and during the following 5 minutes after quenching (see Supplementary Video 1 for a recording of this experiment). Following this second set of experiments, we repeated equibiaxial mechanical testing in 20 cycles to 1000mN of force. See Figure 1B-C for a depiction of the two protocols.

### Histology & Two-photon microscopy

We heat-treated 6mm discoid skin samples in 37°C, 55°C, 75°C, and 95°C 1xPBS for either 10 seconds or 5 minutes and subsequently fixed half of those samples in 10% formalin for 24 hours before shipping them in ethanol to a commercial histology service (Histoserv, Inc., Germantown, MD, United States). Histoserv stained our tissues with Masson’s trichrome and Picrosirius red. Upon receipt of the stained tissue sections we imaged all samples on a 4X optical microscope (BX53, Olympus, Center Valley, PA, United States). The other half of the samples, we imaged under our two-photon microscope using our previously developed imaging protocol in which we excited the tissue under 900nm and epi-collected the emission signal with a 460 ± 25 nm filter to collect collagen’s second harmonic generation signal at a resolution of 1024 by 1024 pixels (over a 500 *µ*m x 500 *µ*m region of interest within the dermis of the skin samples, see Figure 2. Here we followed our own and others’ previous work [24, 25, 26].

**Figure 2:**
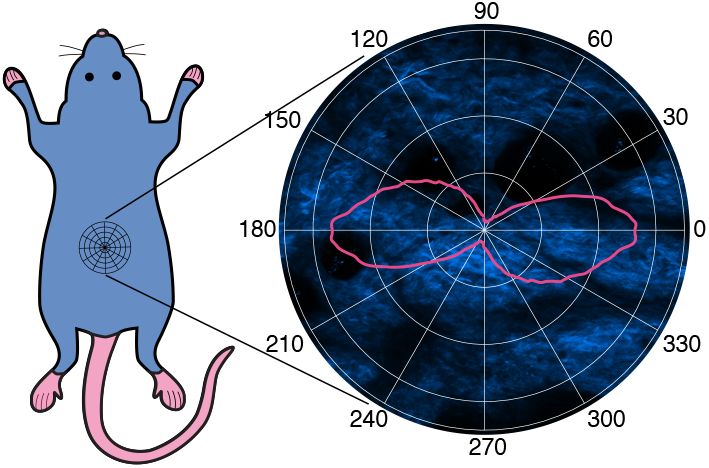
Representative second harmonic generation image of the collagen orientation in dorsal mouse skin. Based on an orientation distribution analysis in ImageJ, we also show the collagen orientation distribution function indicating a clear anisotropy with collagen’s predominant orientation in lateral direction [27]

### Data analysis

First, we evaluated tissue shrinkage based on the photographs taken before and after heat treating in Experiment #1. To this end, we used a custom Matlab program to digitize the perimeter of each sample and automatically quantified area shrinkage as the percentage area change following heat treatment. Additionally, we computed longitudinal and lateral strain as percentage width and height changes, respectively.

Second, we evaluated the biaxial constitutive behavior of skin before and after heat treatment in Experiments #2 and #3. To this end, we evaluated the force strain data during the downstroke of the last biaxial testing cycle. To compute Green-Lagrange strain, we conducted digital image correlation based on the stamp marks on the tissue samples. Next, we transformed force measurements into membrane tension based on the deformed sample width. For each stress-strain curve, we evaluated “toe-stiffness”, i.e., stiffness at small strains, and “calf-stiffness”, i.e., stiffness at large strains by computing the respective tangent moduli.

### Statistics

All data are shown as mean ± standard deviation and sample numbers are provided in each figure caption. We conducted all statistical analyses via linear mixed models as implemented in “R” (Version 4.0.3). Specifically, we used the *afex* library as we previously reported [28].

## Results

We tested a total of 141 skin samples in our isotonic and isometric biaxial tests and our imaging protocols. The results of our isotonic tests are summarized in Figure 3. Overall, we found that skin shrinks when treated with 1xPBS of 75°C and 95°C, but not when treated with 37°C and 55°C 1xPBS, see Figures 3A-C. When samples shrunk, they shrunk more in lateral direction than in cranial direction (p*<*0.001 for both 75°C and 95°C), see 3D. This shrinkage appeared to occur at a faster rate at 95°C than at 75°C, but in both cases equilibrated to similar strains (p=0.4844 and p=0.3870 in cranial and lateral direction, respectively). Thus, upon heat treatment skin shrinks anisotropically and at a rate that is temperature-dependent.

**Figure 3:**
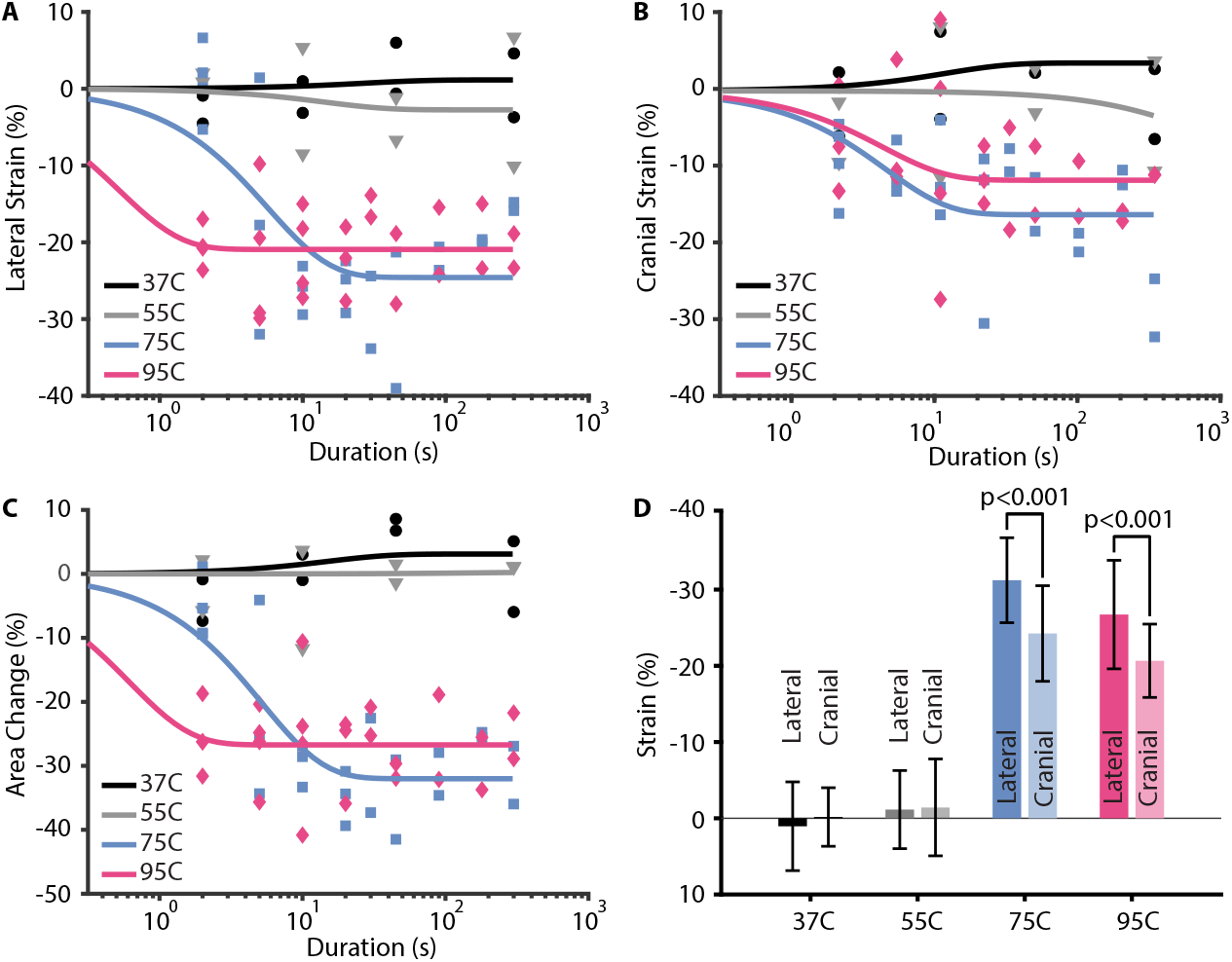
Heat treatment with 75°C and 95°C 1xPBS induces anisotropic shrinkage under isotonic conditions. A) Heat treatment with 37°C (n=9) and 55°C (n=9) did not induce significant lateral strain in discoid skins samples, while treatment with 75°C (n=24) and 95°C (n=26) did. Interestingly, skin shrunk faster at 95°C than at 75°C, but reached similar equilibrium values after 5 minutes. B) Shrinkage in cranial direction mimicked the qualitative shrinkage behavior in lateral direction. However, quantitatively, skin shrunk more in lateral than in cranial direction (see D). C) Heat treatment-induced area shrinkage reflected the qualitative behavior in lateral and cranial directions. D) At equilibrium, 37°C and 55°C treatments did not result in shrinkage, while 75°C and 95°C treatments resulted in significant anisotropic shrinkage, most notably in the lateral direction

In addition to isotonic biaxial tests, we also conducted isometric biaxial tests. Those tests are summarized in Figure 4. Here we found that when constrained at its boundaries, skin produced significant forces upon exposure to 75°C and 95°C 1xPBS that decayed after an initial peak. In contrast, skin did not produce significant forces when exposed to 55°C 1xPBS, Figures 4A-B and Supplementary Figure S1, respectively. Skin’s response to 75°C and 95°C 1xPBS was also anisotropic in that forces where higher in lateral direction than in cranial direction (p=0.0011 and p=0.0003 at 75°C and 95°C, respectively). Additionally, the peak force also increased with temperature (p=0.0064 and p=0.0031 in lateral and cranial directions, respectively), see 4C. The peak force was also reached significantly faster at 95°C than at 75°C (p*<*0.001 for both directions), but did not differ between directions (p=0.6696 and p=0.9158 for 75°C and 95°C, respectively), see Supplementary Figure S2. Additionally, the time constant with which the force decayed after reaching its peak was significantly shorter during the 95°C treatment than after the 75°C treatment (p*<*0.001 for both temperatures), see Figure 4D. The decay time, like the time to peak, did not differ with direction (p=0.6934 and p=0.9379 for 75°C and 95°C, respectively). Thus, upon heat treatment constrained skin produces significant anisotropic forces at rise and decay times that are temperature-dependent. Note, after five minutes of exposure to 75°C and 95°C, these samples could not be tested mechanically. They had softened to the point where stretching them resulted in tear-outs from our fixtures. In contrast, five minutes of exposure to 55°C rendered the samples testable and only marginally affected skin’s mechanics (see Supplementary Figure S3).

**Figure 4:**
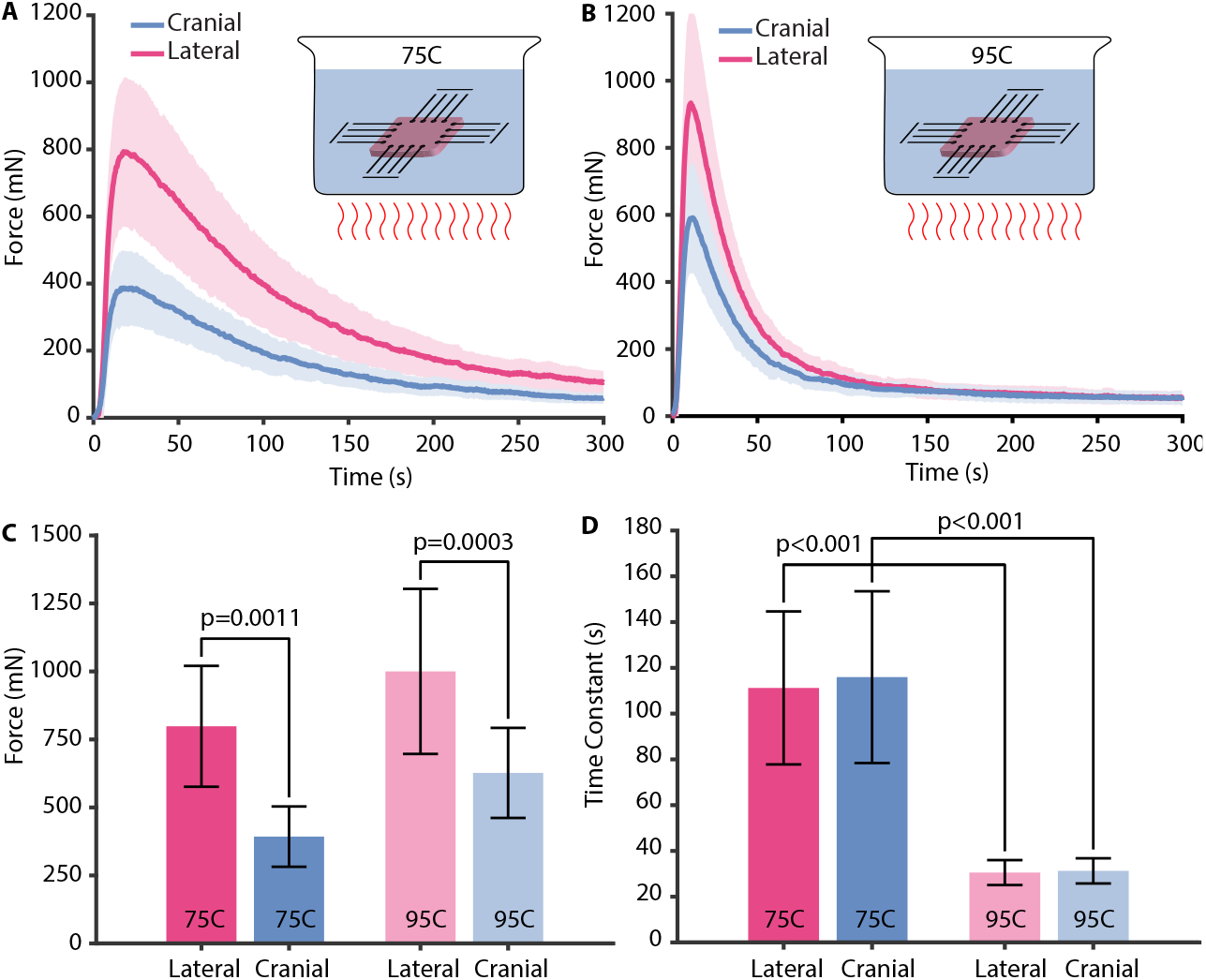
Heat treatment with 75°C and 95°C 1xPBS induces anisotropic forces under isometric conditions. A) In both lateral and cranial direction, heat treatment with 75°C (n=9) 1xPBS induced forces that rose quickly before exponentially decaying to almost zero within 5 minutes. Forces were higher in lateral than cranial direction (see C). B) Heat treatment with 95°C (n=9) 1xPBS qualitatively mimicked the response to 75°C. C) Heat treatment with 95°C reached higher peak forces than with 75°C (note, for clarity we omitted the lines indicating this difference: p=0.0064 and p=0.0031 in lateral and cranial directions, respectively) and forces were higher in lateral direction than cranial direction for both 75°C and 95°C. D) Additionally, peak forces in response to 95°C heat treatment decayed faster

Thus, in another set of experiments we exposed skin to 90°C 1xPBS, but interrupted the force decay by quenching (i.e., quickly cooling with 37°C 1xPBS) samples after exposure times between 0 and 30 seconds. Upon cooling, forces remained approximately constant rather than continuing to decay indicating that the internal damage processes had been interrupted, see Supplementary Video 1. After quenching, we tested those samples biaxially - in addition to the biaxial tests we conducted before the heat treatment. A representative biaxial data set before and after 10 seconds of heat treatment (on the same sample) anecdotally demonstrates that heat treatment stiffened skin at low strains, but reduced its stiffness at high strain, see Figure 5A. Interestingly, we found that stiffening and softening at low and high strains, respectively, was not clearly correlated to the exposure times in the interval between 0 and 30 seconds, see Figure 5B. When grouping the data - in other words, ignoring exposure time - we could demonstrate a statistically significant skin softening after heat treatment at high strains (p*<*0.001 in both lateral and cranial direction), but failed to show statistically significant stiffening after heat treatment at low strains (p=0.061 and p=0.5348 in lateral and cranial direction, respectively), see Figures 5C-D. Thus, skin does soften at high strains and may stiffen at low strains when treated with 90°C 1xPBS, where the latter phenomenon remains to be demonstrated.

To identify the source of the heat-induced changes to skin illustrated in Figures 3-5, we investigated (micro-)structural changes in our skin samples using linear and nonlinear optical methods. Specifically, we stained samples after isotonic heat exposure to 37°C, 5°C, 75°C, and 95°C with picrosirius red (and for reference with Masson trichrome). We found that the (birefringent) picrosirius red signal remained strong after exposure to 37°C and 55°C, but vanished after 5 minutes of 75°C exposure and 10 seconds as well as as 5 minutes of 95°C exposure, see Figure 6. Thus, heat treatment of skin removes the source of birefringence in a time- and temperature-dependent manner. Similarly, we imaged skin samples after the same isotonic heat treatment in our two-photon microscope and collected its SHG signal. We found that the SHG signal remains strong after exposure to 37°C and 55°C, but vanished after 5 minutes of 75°C exposure and 10 seconds as well as 5 minutes of 95°C exposure, see Figure 7. Thus, heat treatment of skin also removes the source of SHG in a time- and temperature-dependent manner and supports our histological findings.

**Figure 5:**
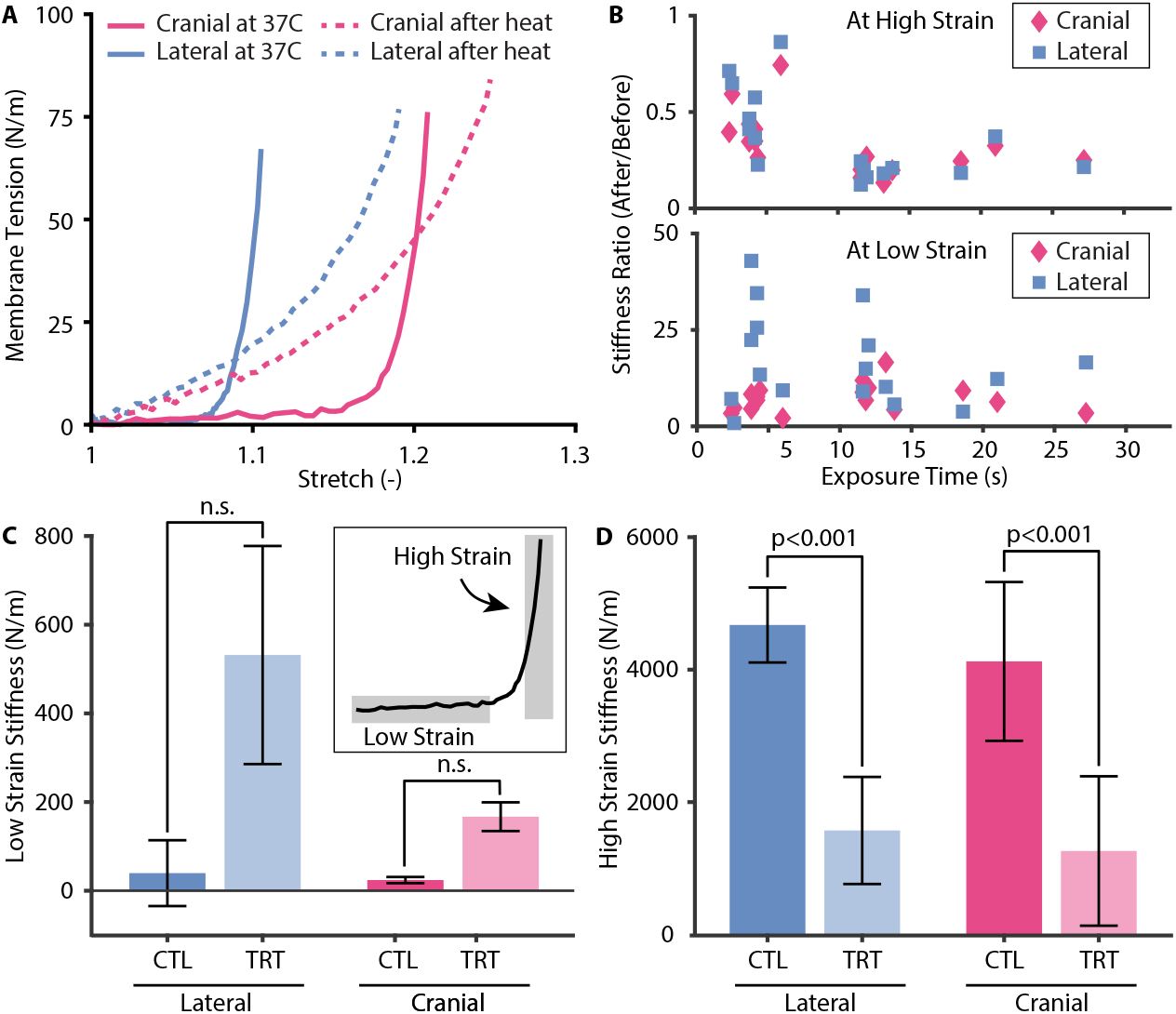
Heat treatment with 90°C 1xPBS alters constitutive behavior of skin. A) Example raw data of biaxial test before and after heat treatment for 10 seconds. B) The relationship of heat treatment duration and stiffness increase (at low strain) or decrease (at high strain) was subject to large variations and appeared uncorrelated (n=17). C) When grouped, heat treatment-induced stiffening of skin at low strain, and D) heat treatment-induced reduction in stiffness of skin at high strain

**Figure 6:**
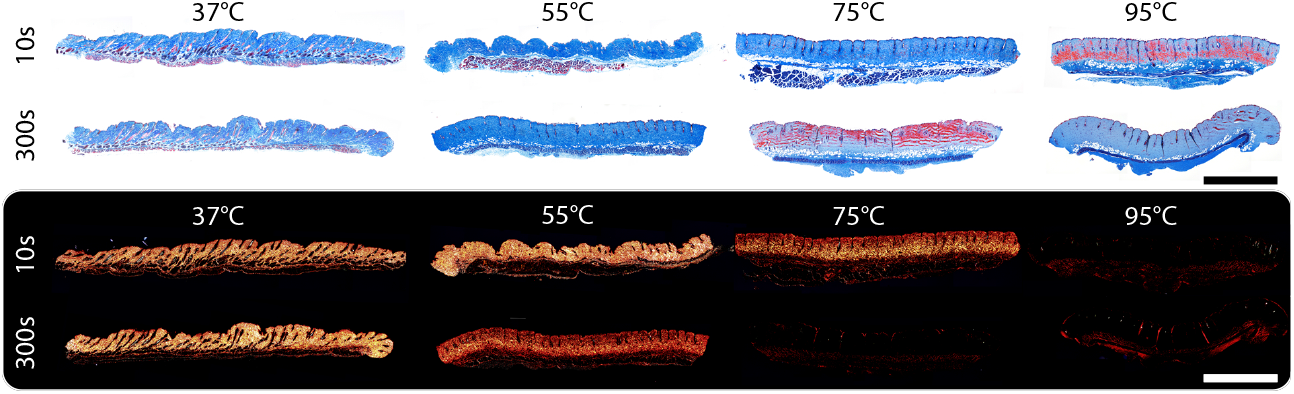
Heat treatment with 75°C and 95°C 1xPBS denatures collagen as demonstrated by loss of birefringence. This effect is time- and temperature-dependent as demonstrated by residual birefringence after 10s of treatment with 75°C, but loss of birefringence after 5 minutes, while heat treatment with 95°C resulted in loss of birefringence at 10s and 5 minutes. We confirmed each time-temperature-pair with an independent set of samples (not shown). Scale bar = 1mm

**Figure 7:**
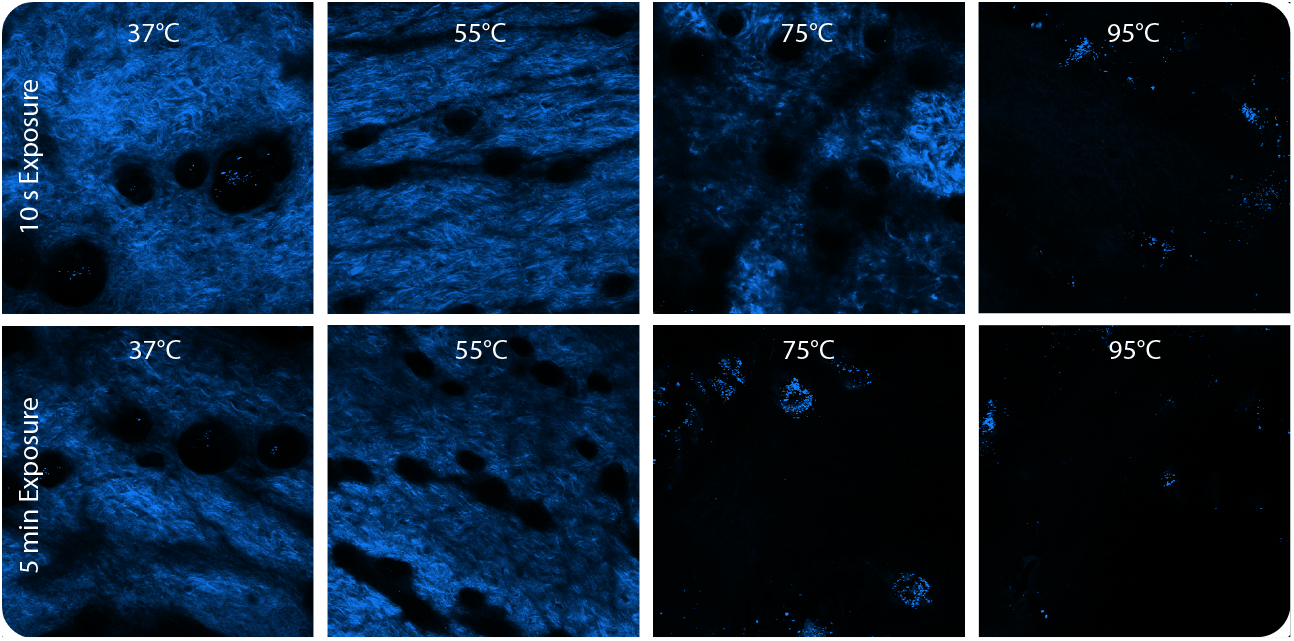
Heat treatment with 75°C and 95°C 1xPBS denatures collagen as demonstrated by loss of second harmonic generation (SHG) signal. This effect is time- and temperature-dependent as demonstrated by residual SHG signal after 10 seconds of treatment with 75°C, but loss of SHG signal after 5 minutes, while heat treatment with 95°C resulted in loss of SHG signal at 10 seconds and 5 minutes. Note, we confirmed each time-temperature-pair with an independent set of samples (not shown). Image size 500*µ*m x 500*µ*m

## Discussion

In our current work, we investigated the biaxial mechanics of skin after treatment with 37°C, 55°C, 75°C, and 95°C. Our work was motivated by missing information on how skin’s anisotropic microstructure and mechanical constitutive behavior impacts its thermo-mechanical response to heat. We designed our experiments to investigate the thermo-mechanical response of skin to both isotonic loading - that is, traction-free - and isometric loading - that is, with fixed boundaries - in a temperature- and time-dependent manner. Our findings confirmed many previous studies on the native, i.e., not heated, anisotropic microstructure and mechanical constitutive behavior of skin [29]. Additionally, our results were consistent with non-directionally-dependent previous studies on skin’s thermo-mechanical response to heat treatment [22, 23]. In addition to confirming previous findings, we unveiled significant, anisotropic thermo-mechanical effects in skin’s response to heat treatment. We also used linear and nonlinear microscopy to confirm that the thermo-mechanical response of our samples was not temporary but followed from molecular damage.

### Summary of findings

Our main findings were that heat treatment leads to significant skin shrinkage under isotonic conditions and produces significant forces under isometric conditions. Under isometric conditions, skin also softens at large strains. These effects are temperature-, time-, and direction-dependent. Specifically, our findings on temperature-dependence are: i) Under isotonic conditions, heat treatment induces skin shrinkage at temperatures larger than 55°C; ii) Under isotonic conditions, higher temperatures induce faster shrinkage, but do not lead to more shrinkage; iii) Under isometric conditions, heat treatment induces forces at temperatures larger than 55°C; iv) Under isometric conditions, higher temperature induces forces faster, lead to larger forces that also decay faster. Our findings on time-dependence are: i) Under isotonic conditions, shrinkage sets in within seconds but then equilibriates; ii) Under isometric conditions, force production peaks within seconds, but then continuously decays until dissipated. Finally, our findings on direction-dependence are: i) Under isotonic conditions, heat treatment (*>* 55°C) induces more shrinkage in lateral direction than in cranial direction; ii) Under isometric conditions, heat treatment (*>* 55°C) induces larger forces in lateral direction than in cranial direction; iii) Under isometric conditions, heat treatment (*>* 55°C) reduces mechanical anisotropy.

Based on our histological and two-photon based analysis, skin’s thermo-mechanical response to heat treatment was correlated with a degradation of optical properties – birefringence and SHG – that are related to collagen’s structure [30, 31]. In other words, it is, at least in part, the heat treatment-induced denaturing of collagen that leads to above thermo-mechanical phenomena.

### Hypothetical model of skin’s thermo-mechanical response to heat treatment

Combining our thermo-mechanical and optical observation, we suggest the following hypothetical model for the effects of heat treatment on skin: Thermal destruction of intra- and inter-molecular bonds in the collagen molecule lead to the disruption of collagen’s rod-like alpha helix [32]. Without these bonds, coiling results in an energetically favorable state and thus leads to tissue shrinkage in the absence of external forces [33]. Phenomenologically, this transition may be described as redefinition of the stress-free reference state of the collagen molecule to shorter lengths [34]. The increased density of collagen in the lateral direction in comparison to the cranial direction amplifies this effect in a directionally-dependent manner [35]. The thermal destruction of the intra- and inter molecular bonds is likely driven by an Arrhenius-type, time-dependent mechanism, thus explaining the increased rate of shrinkage at higher temperatures [5]. In the presence of constraints, the same thermal destruction of intra- and inter-molecular bonds occurs. However, external tractions prevent coiling of the collagen molecular, thus, leading to the production of forces as the stress-free reference configuration of the collagen molecule shifts to shorter lengths. Here, again, anisotropic distributions of collagen result in a direction-dependent number of coiling collagen fibers and thus a higher production of force in the lateral direction than the cranial direction. Also, here, again, Arrhenius-type reactions lead to a temperature dependent production of these forces. More mysterious, however, is the specific temporal evolution of the force production. If the forces were produced through the constrained coiling of collagen, why do they decay over time? We propose two alternative models. First, we propose that continued exposure to heat may additionally degrade collagen. In turn, collagen may entered a third, further degraded state (beyond being coiled). Second, and alternatively, it may be that, under force, the denatured collagen molecule may undergo viscoelastic stress-relaxation. In either case, time-dependence of the additional denaturation or, alternatively, viscoelastic stress-relaxation, leads to faster decay times at higher temperatures. This hypothetical model also explains our observation that heat treated skin anecdotally stiffens at low strains and softens at high strains. Under native, pre-treated conditions, wavy collagen fibers are activated only at large strains, a mechanism that lends skin – and many other collageneous soft tissues – their classic J-shaped or strain stiffening stress-strain behavior [36]. Upon heat treatment, the reduced reference configuration of collagen leads in our hypothetical model to an early activation of its contribution to the tissue’s resistance to external loading and thus a stiffening at small strains. On the other hand, the destruction of inter- and intra-molecular bonds reduces collagen’s load-bearing capacity and leads to a softening at large strains.

### Shortcomings, unexplained phenomena, and future work

We believe that our work is an important step toward understanding the biaxial mechanics of thermally denaturing skin. However, our work is naturally subject to limitations and there is a great need for additional work. For example, Chen et al have shown that the thermo-mechanics of collageneous tissues are not only time-, and temperature-dependent, but also load-dependent [7]. This additional determinant of skin’s response to thermal treatment should, in the future, be added to our observed dependence on time, temperature, and direction. From an experimental design perspective, we observed the mechanical manifestations to thermal treatment – shrinkage, force production, and change in constitutive behavior - and the microstructural origins - denaturation of collagen - independently. Future studies should aim at a design that simultaneously images skin’s microstructure while recording its mechanical response to heat treatment [37]. Additionally, we failed to demonstrate a statistical relationship between tissue stiffening and softening – at low and high strains, respectively – and heat exposure times. Our failure to do so is most likely related to the high-rate, nonlinear temperature response curve as seen in the time evolution of force production. Future experiments toward establishing a clear relationship between skin stiffening/softening and exposure times will have to take great care to control exposure times more accurately than we were able to do in this present study. Finally, and most importantly, we have posed a theoretical model connecting skin’s thermo-mechanical response to heat treatment to its microstructural origins. As a first step toward confirming this model, we will implement this model numerically and test its ability to predict our observations.

## Conclusion

We conclude that thermal treatment of skin above 55°C results in time-, temperature-, and direction-dependent shrinkage under isotonic conditions. The largest shrinkage occurs in the direction of the predominant collagen orientation. Similarly, thermal treatment of skin above 55°C results in time-, temperature-, and direction-dependent force production under isometric conditions. The largest forces occur in the direction of the predominant collagen orientation. Additionally, we conclude that thermal treatment of skin results in softening of skin at high strains. These thermo-mechanical phenomena are driven by the denaturation of collagen as demonstrated through histology and two-photon microscopy. This work is a critical step toward developing a fundamental understanding of the relationship between skin microstructure, mechanics, and its response to thermal treatment. Ultimately, this knowledge will be critical to optimally treating burn wounds, designing medical devices, and optimizing aesthetic modifications.

## Acknowledgements

We acknowledge the National Science Foundation for their partial support of this project via Grant #1916663 (Rausch, Buganza), the National Institutes of Health for their partial support of this project via Grant F31HL145976 (Meador), and the University of Texas at Austin Graduate School Continuing Student Fellowship (Sugerman).

