## Supplementary figures and images for "The biaxial mechanics of thermally denaturing skin - Part I: Experiments"

### Supplementary Figure S1

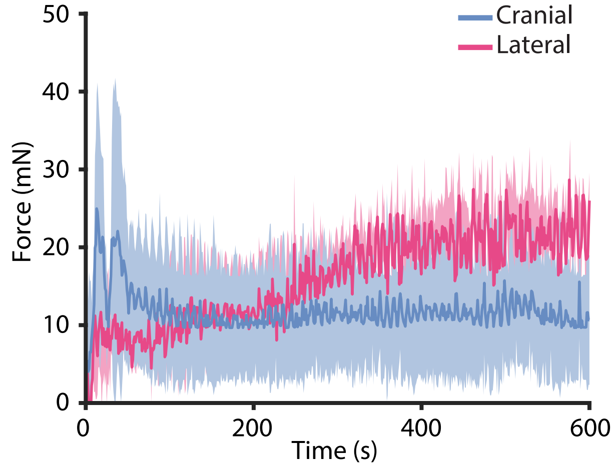

### Supplementary Figure S2

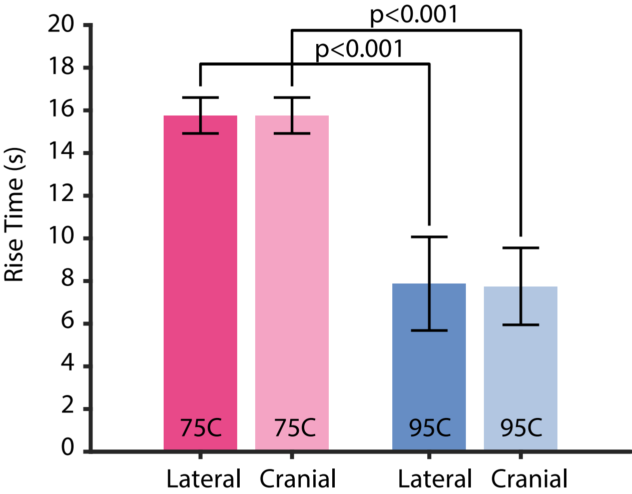

### Supplementary Figure S3

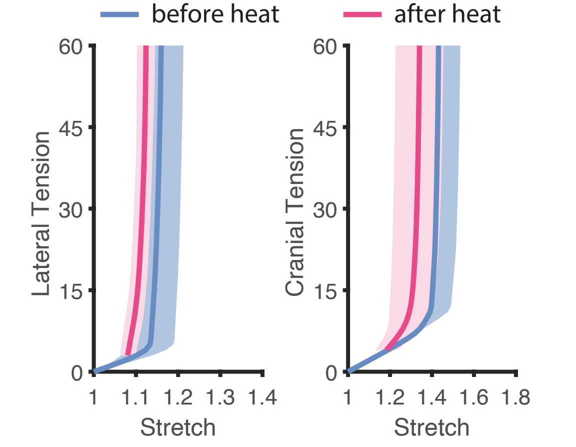

### Supplementary Video 1

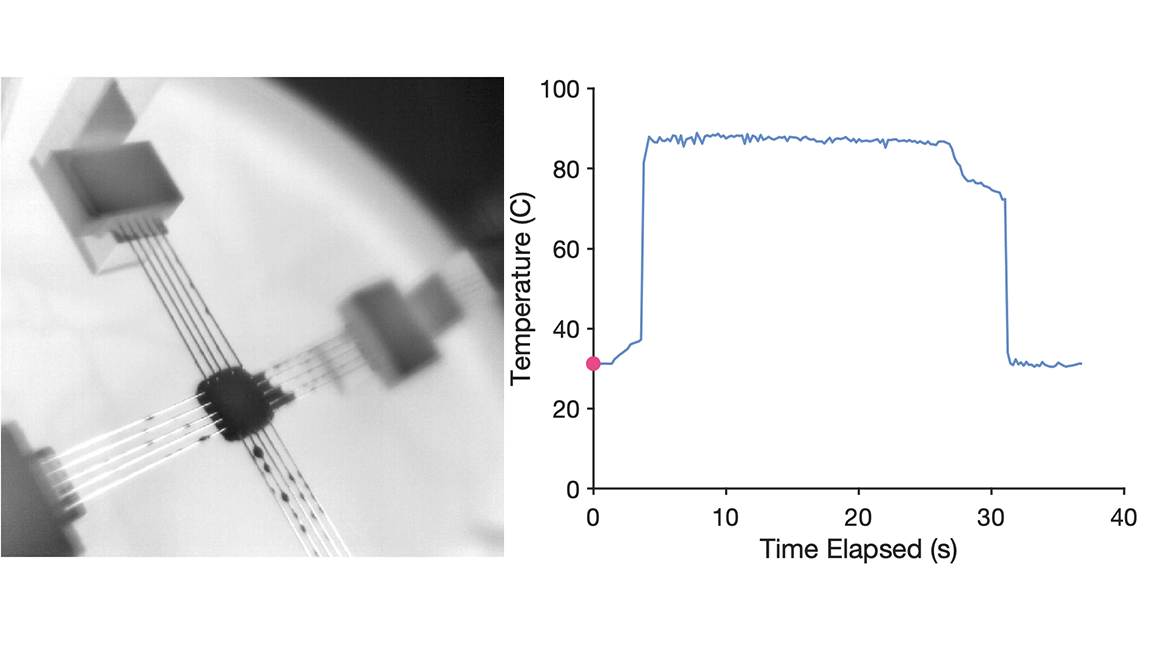
